# multiTFA: a Python package for multi-variate Thermodynamics-based Flux Analysis

**DOI:** 10.1101/2020.12.01.407387

**Authors:** Vishnuvardhan Mahamkali, Tim McCubbin, Moritz Emanuel Beber, Esteban Marcellin, Lars Keld Nielsen

**Affiliations:** Australian Institute for Bioengineering and Nanotechnology (AIBN), The University of Queensland, 4072 Brisbane, Australia; The Novo Nordisk Foundation Center for Biosustainability, Technical University of Denmark, 2800 Kgs. Lyngby, Denmark

## Abstract

**Summary:** We achieve a significant improvement in thermodynamic-based flux analysis (TFA) by introducing multivariate treatment of thermodynamic variables and leveraging component contribution, the state-of-the-art implementation of the group contribution methodology. Overall, the method greatly reduces the uncertainty of thermodynamic variables.

**Results:** We present multiTFA, a Python implementation of our framework. We evaluated our application using the core *E. coli* model and achieved a median reduction of 6.8 kJ/mol in reaction Gibbs free energy ranges, while three out of 12 reactions in glycolysis changed from reversible to irreversible.

**Availability and implementation:** Our framework along with documentation is available on https://github.com/biosustain/multitfa.

## 1 Introduction

Constraint-based analysis of metabolic network models is used widely to explore metabolic phenotypes and guide metabolic designs (O’Brien *et al*., 2015). Thermodynamic-based flux analysis (TFA) imposes thermodynamic constraints on constraint-based models, in order to obtain thermodynamically valid metabolic fluxes and metabolite concentration profiles (Henry *et al*., 2007). TFA provides an ideal mechanism for incorporating metabolomics data into genome-scale modelling. TFA is also a critical pre-processing step when performing sampling based fitting and exploration of large kinetic models (Saa and Nielsen, 2017).

Thermodynamic constraints rely on the calculation of Gibbs free energies of compounds and reactions. The current best method for estimating standard Gibbs free energy of reaction (Δ_r_G^’^_i_) uses the component contribution method, which combines reactant and group contribution methods while maintaining thermodynamic consistency (Noor *et al*., 2013; Flamholz *et al*., 2012).

Accommodating the errors in the estimated Δ_r_G^’^_i_ presents a challenge. We cannot introduce independent slack in each Δ_r_G^’^_i_, since this would cause inconsistent thermodynamics with non-zero Gibbs energy loops. The original TFA implementation was based solely on the group contribution method (Henry *et al*., 2007). It avoided inconsistency by computing Δ_r_G^’^_i_ within the algorithm from ‘groups’ treated as independent variables allowed to vary within their individual 95% confidence intervals, i.e., approximately two standard deviations (*SD*) around the mean 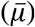 (Henry *et al*., 2007). More recently, the loop issue has been addressed by using metabolite formation energies rather than reaction energies in pyTFA (Salvy *et al*., 2019). These formation energies can be user defined or calculated as a linear combination of respective group Gibbs free energies (from a suitable database). The pyTFA algorithm also treats formation energies as independent variables which are allowed to vary in the range 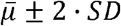 (Salvy *et al*., 2019).

It is not optimal to use the n-box formed from individual 95%-confidence intervals to capture the range of feasible values in a multivariate distribution, such as the full set of formation energies generated by the component contribution method. Firstly, the n-box does not define a 95%-confidence range for the mean vector of formation energies. Secondly, and more importantly, it does not capture the correlation in the distribution. This is particularly problematic using formation energies for substrates and products linked through a reaction, since they will tend to be highly correlated. For illustration consider the multivariate normal distribution estimates for ATP and ADP with the following mean vector and covariance (Σ) matrix (see details later):

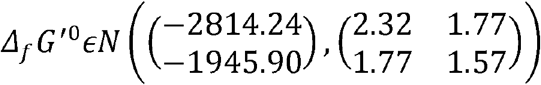

Unsurprisingly, the estimates are highly correlated (0.928), which is reflected in the cigar shaped 95%-confidence ellipse (Fig. 1A, blue line). While the n-box (red box) almost captures the range, it fails to capture the correlation. Since ATP and ADP are commonly found on either side of a reaction, the most important value is the difference in free energy. The range for the difference is much smaller using the proper confidence region (2.9 kJ/mol) compared to using the n-box (10.9 kJ/mol). Using multivariate confidence regions effectively ensures that we cancel out common error contributions on either side of a reaction (Haraldsdóttir *et al*., 2012). We note that the original method from Henry *et al*. achieved error cancellation for common groups but did not address correlation between group estimates.

**Figure 1.**
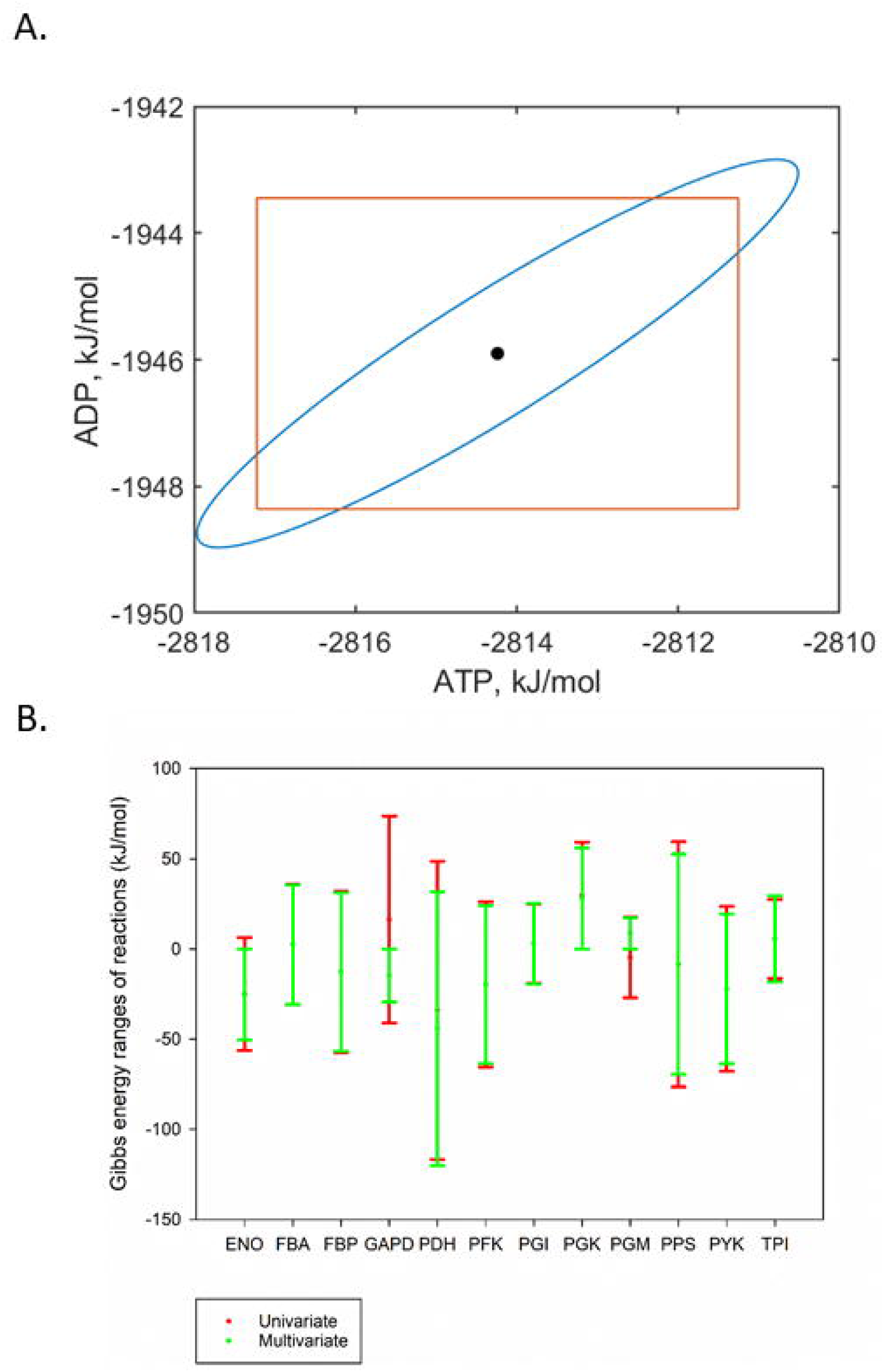
A) Comparing the 95% confidence ellipse (blue) to the n-box (orange) for the eQuilibrator estimates of ATP and ADP formation energies. B) Comparison of the Gibbs free energy of reaction ranges across the glycolytic pathway estimated using the n-box (univariate) and the multiTFA (multivariate) methods. Three reactions change from reversible to irreversible when using a multivariate treatment (ENO, GAPD, and PGM).

multiTFA is an internally consistent TFA framework with multivariate treatment of errors in formation energies. Constraining formation energies within the 95% confidence ellipsoid rather than the n-box more accurately captures the range of values, while narrowing the likely range of free energies of reaction and concentration values.

## 2 Materials and Methods

The constraints in TFA are (Salvy *et al*., 2019)

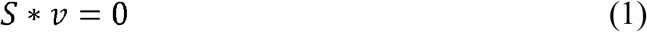

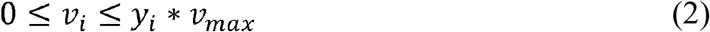

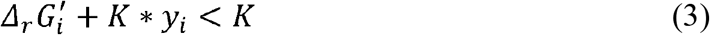

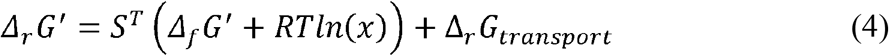

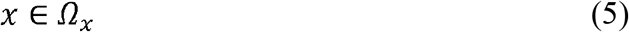

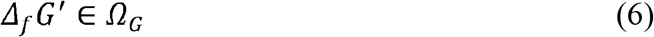

where *S* and *v* are the stoichiometric matrix and flux vector, respectively. Reactions only progress in the forward direction (i.e., reversible reactions are split in two), and only if the binary coupling variable, y_i_, is 1 (2), which can only happen when the Gibbs free energy of the reaction (*Δ_r_G*′)is negative (3) (K is a large positive constant).

*Δ*_*r*_*G′* is calculated from the formation energies (*Δ_f_G*′) and concentrations (x) of the metabolites (4). For transporters, the Gibbs free energy of transport was calculated as detailed in (Jol *et al*., 2010). Our implementation automatically detects transporters and predicts the species that is being transported based on the pKa value and the compartment pH. Users are also able to explicitly define the charged form of the transported metabolite and transportation mechanism. For the calculation of Gibbs free energies of reaction at non-standard conditions, users can input the range for each metabolite concentration, *Ω_x_*(5). Where not specified, metabolites can assume pre-defined compartment specific bounds or otherwise adopt loose bounds (10^−5^ – 10^−2^ M).

The formation energies are estimated using the component contribution method (Noor *et al*., 2013) and adjusted for compartment specific pH and ionic strength (Alberty, 2005; Haraldsdóttir *et al*., 2012). It is assumed that the estimate follows a multivariate normal distribution, *Δ*_*f*_*G*′*∈N*(*µ,Σ*).TFA allows for noise in the *Δ*_*f*_*G*′estimate by defining a region, *Ω_G_* (6). A common approach is to use as *Ω_G_*the n-box defined by the individual 95%-confidence intervals for each formation Gibbs energy, i.e 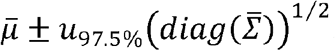. This is not a true 95% confidence region for the multivariate estimate: it greatly underestimates the range of individual variables and ignores the correlation between related compounds such as ADP and ATP. A more appropriate region, *Ω_G_*, would be the 95%-confidence ellipsoid defined by:

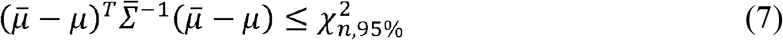

Introducing this constraint converts the problem from a Mixed Integer Linear Problem (MILP) to a Mixed Integer Quadratic Constraint Problem (MIQCP).

In general, *Σ* does not have full rank and (7) cannot be used directly. *Δ_f_G*′ is calculated using the component contribution method as

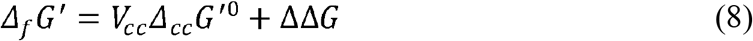

Where *V*_*cc*_ is the metabolite component composition vector, Δ_*cc*_*G*^′0^ is a vector of component and group standard Gibbs energies, and ΔΔ*G* is a (deterministic) adjustment for compartment pH, pI and Mg concentration. This estimate is the fit of the component contribution model to the thermodynamic reference data and is assumed to follow a multivariate normal distribution, *Δ_cc_G*^′0^*ϵN* (*µ*_*cc*_, *Σ*_*cc*_). We can express this distribution as

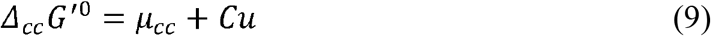

where *CC*^*′*^ = *Σ*_*cc*_ and *uϵN* (0, *I*). Allowing for *Σ*_*cc*_ not having full rank, we use LDL decomposition to achieve pivoted Cholesky decomposition finding 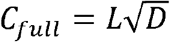. *C*_*full*_ has *n = rank*(*Σ*_*cc*_)non-zero columns and we obtain *C* by removing the remaining columns. Finally, we define a 95%-confidence circle for the n-dimensional standard normal distribution

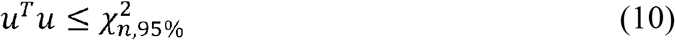

Using (10), we still need to solve an MIQCP, but this is numerically more robust.

We compared multiTFA with the n-box approach using an *E. coli* core model (e_coli_core) (Orth *et al*., 2010). We determined the Gibbs free energy ranges using either (a) the “conventional” 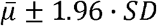 n-box or (b) a multivariate treatment of the errors in the formation energy estimate used in multiTFA. Despite the n-box covering far less than the 95%-confidence range for individual formation energies, the estimated ranges for the Gibbs free energy of reaction were broader than using the confidence ellipsoid. The median reduction in Gibbs free energy ranges was 6.8 kJ/mol (Figure 1b; Supplementary Data), highlighting the significant potential for error cancellation between compounds captured in the correlation matrix. The reduction in Gibbs free energy ranges is reflected in a reduction in the reaction flux ranges (Supplementary Data).

### 2.1 Usage & Implementation

The Python package comes with example scripts to demonstrate the usage of different functionalities. The software takes a typical COBRA model as input and generates a MILP Optlang object (Jensen *et al*., 2017) for the n-box approach that can be directly solved with COBRApy (Ebrahim *et al*., 2013). If the user has Gurobi or CPLEX solver installed, the software will generate solver specific MIQCP objects to solve the multiTFA problem. For users without Gurobi or CPLEX, an alternative implementation of multiTFA is provided that uses random sampling of the surface of the confidence ellipsoid and a MILP solver to determine the maximum range. The exit criterion of the sampler can be chosen as either (1) the number of samples since last improvement or (2) a fixed number of samples followed by use of a generalized extreme value distribution to infer the maximum value.

The implementation is available at https://github.com/biosustain/multitfa. The framework is currently compatible with models that use different identifiers (SEED, KEGG, BIGG among others) for matching metabolite information against the thermodynamic database. We use the eQuilibrator API to retrieve data matrices for calculating the formation energies and covariance matrix (Noor *et al*., 2013).

## 3 Conclusion

Using a multivariate confidence ellipsoid to describe the feasible range in the Gibbs free energy of formation estimate, multiTFA is able to account for a more realistic (and broader) range in individual estimates of formation energy, while simultaneously using correlation to reduce the ranges for the derived Gibbs free energies of reactions.

